# Local cytokine changes in response to mesenchymal stem cell-derived extracellular vesicles-based therapy in rat spinal cord injury

**DOI:** 10.1101/2024.10.07.616981

**Authors:** Ilyas Kabdesh, Ekaterina Garanina, Alexander Kostennikov, Albert Rizvanov, Yana Mukhamedshina

## Abstract

**Aim:** To investigate the effects of mesenchymal stem cell-derived extracellular vesicles (MSC-EVs), encapsulated in a fibrin matrix (FM), on pro-inflammatory and anti-inflammatory cytokine levels in a rat model of spinal cord injury (SCI) 60 days post-injury.

**Methods:** MSCs were isolated from rat adipose tissue and cultured to obtain MSC-EVs using cytochalasin B. MSC-EVs encapsulated in the FM were applied to the injury site at doses of 5 and 10 µg. Four experimental groups included SCI without treatment, SCI with FM application, and SCI with FM+EVs at 5 and 10 µg doses. A multiplex assay was conducted to measure 23 cytokines in spinal cord tissue homogenates.

**Results:** FM application to the injury site exhibited both pro- and anti-inflammatory cytokine shifts, with the most pronounced effect for G-CSF (2.8-fold) potentially due to the hemostatic properties of the FM. MSC-EVs led to significant modulation of cytokine levels. In the SCI FM+EVs10 group, concentrations of IL-10, IL-1b, IL-5, and IL-17A were significantly lower compared to the SCI and SCI FM groups. Here the most pronounced change was observed for IL-10, which decreased by 2.4-fold.

**Conclusion:** Combined treatment with MSC-EVs and FM significantly influenced pro- and anti-inflammatory cytokine levels, demonstrating a dose-dependent effect. These findings highlight the potential of EVs to modulate inflammatory responses and promote regeneration in the chronic phase of spinal cord injury.

## Introduction

Spinal cord injury (SCI) is one of the most serious public health concerns, as it often results in permanent disability. Primary mechanical damage to nerve tissue triggers a cascade of secondary pathological reactions, with neuroinflammation playing a key role. A crucial component of this process is pro-inflammatory cytokines, whose excessive production exacerbates neural tissue damage. Therefore, approaches to post-traumatic spinal cord repair should incorporate anti-inflammatory effects.

The use of mesenchymal stem cells (MSCs) and/or their paracrine mediators encapsulated in extracellular vesicles (MSC-EVs) is considered a promising therapeutic strategy. Studies using a rat model of SCI have shown that systemic administration of MSC-EVs reduces the levels of pro-inflammatory cytokines such as TNF-α and IL-1β, while increasing the production of anti-inflammatory molecules like IL-10. In this context, miRNAs, such as miR-21-5p, which regulate inflammation and apoptosis via the FasL signaling pathway, have gained significant research attention. Inhibition of miR-21-5p by MSC-EVs attenuated inflammatory responses, suggesting an important role for miRNAs in neuroprotective mechanisms during SCI [1]. Additionally, beyond apoptosis and neuroinflammation, systemic administration of MSC-EVs has also been shown to modulate angiogenesis, thereby promoting tissue healing by accelerating the elimination of inflammatory mediators and attracting immune cells to the injury site [2].

A study by Romanelli *et al*. (2019) examined the long-term effects of EVs administered intravenously in a rat model of spinal contusion. The study found that during the chronic phase of SCI, both the experimental and control groups continued to lose neural tissue. However, by the end of the experiment, significantly more tissue was preserved in the group receiving intravenously injected EVs compared to the control group [3]. In a follow-up study [4], intraspinal administration of EVs led to a more pronounced suppression of inflammatory responses, a reduction in pro-inflammatory cytokine release (TNF-α and IL-6), and an increase in anti-inflammatory cytokine production (IL-10). Additionally, this approach reduced scar tissue formation and improved motor function compared to systemic EV administration. This difference may be due to localized administration providing a more rapid and targeted effect on inflammatory processes at the injury site [4]. Despite being more invasive than intravenous injection, local delivery of cells or EVs near the spinal cord lesion has generally been shown to be safe when performed slowly and carefully in patients with spinal cord injury [5].

In view of the above, it can be assumed that the local application of MSC-EVs may serve as an effective anti-inflammatory agent in SCI due to its targeted effect on the site of inflammation and reduction of systemic side effects. However, the use of MSC-EVs requires the development of an optimal delivery system that ensures prolonged and controlled release of EVs at the injury site. One promising approach is the use of a fibrin matrix (FM), which is widely employed in clinical practice due to its biocompatibility and natural biodegradability.

Our previous study demonstrated that the application of MSC-EVs promoted the preservation of mature oligodendrocytes and improved functional outcomes in rats in SCI [6]. However, to date, no studies have thoroughly evaluated the effects of combining MSC-EVs with FM on both pro-inflammatory and anti-inflammatory cytokine levels in SCI. The aim of this work was to investigate the impact of co-applying MSC-EVs with FM on cytokine levels in the rat spinal cord during the chronic phase of SCI.

## Materials and methods

All studies were designed to minimize the number of animals and the severity of the procedures. The design of the experiment is summarized in Figure 1.

**Figure 1.**
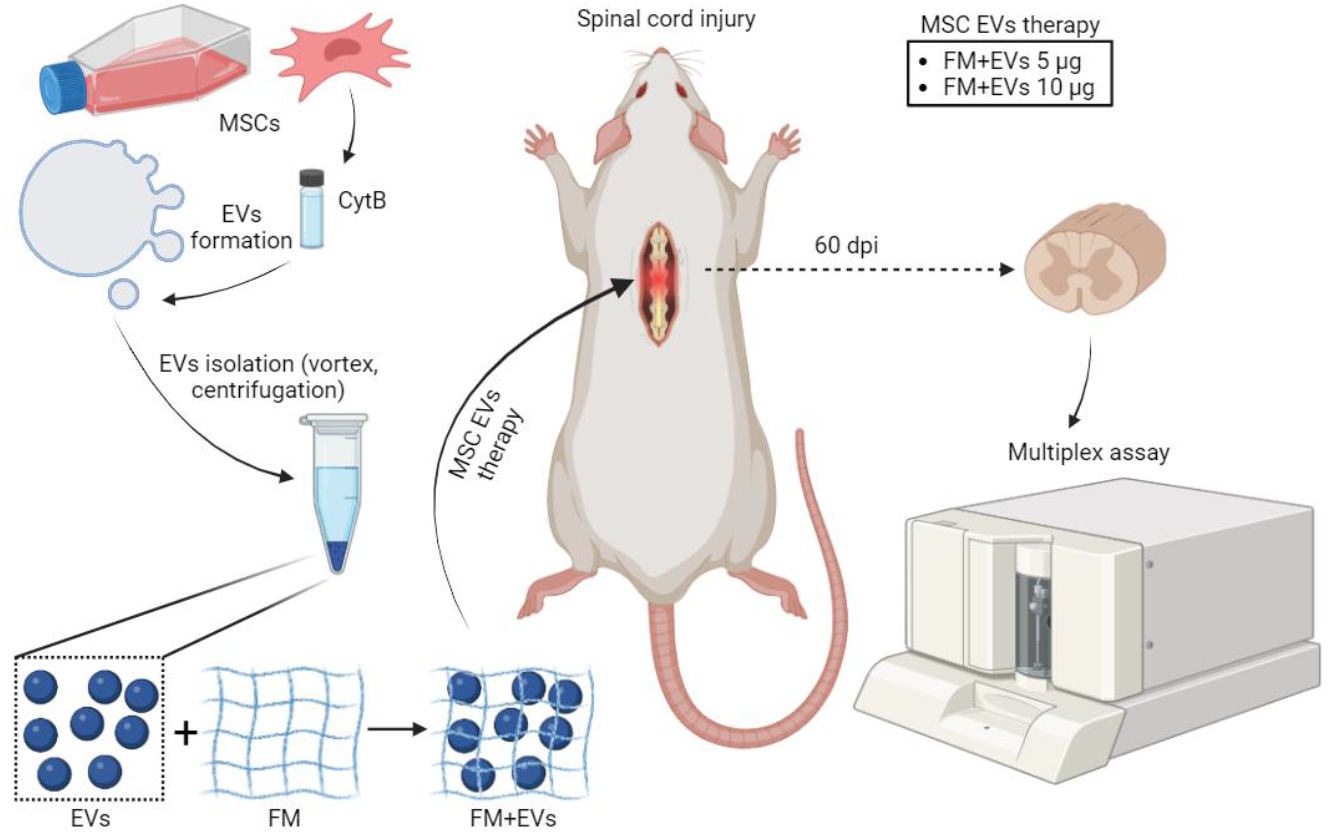
Experimental design (created with Biorender.com). Mesenchymal stem cells (MSCs) were isolated from rat adipose tissue and cultured. Extracellular vesicles (EVs) were isolated from MSCs using cytochalasin B (CytB). EVs were then encapsulated in a fibrin matrix (FM) at doses of 5 µg and 10 µg and applied to the site of spinal cord injury in rats. 60 days post-injury (dpi), the spinal cord was harvested, and multiplex analysis was performed to assess cytokine levels.

### Isolation and Cultivation of Mesenchymal Stem Cells

Mesenchymal stem cells (MSCs) were sourced from the adipose tissue of female Wistar rats weighing 250–300 g (Pushchino Laboratory, Russia). The rats were anesthetized using isoflurane (1.3%, Laboratories Karizoo, Spain) and Zoletil (20 mg/kg, Virbac, France) before undergoing surgery. Adipose tissue was carefully collected in a sterile environment and placed into a container with 0.9% NaCl. It was then processed by homogenizing, followed by a centrifugation rinse in 0.9% NaCl, and finely crushed for 10 minutes at 1500 rpm. Afterward, it was treated with a 0.5% collagenase solution derived from crab pancreas (Biolot, Russia) at 37°C for 1 hour, with constant shaking at 180 rpm. Following this, the homogenate was centrifuged at 1400 rpm for 5 minutes to remove the enzyme solution. The remaining cells were washed in Dulbecco’s phosphate-buffered saline (DPBS, PanEco, Russia), centrifuged again to remove residual enzymes, and prepared for culture. The cells were then grown in DMEM, enriched with 10% fetal bovine serum (FBS), 2 mM L-glutamine, 100 μg/mL streptomycin, and 100 U/mL penicillin (all from PanEco, Russia). The culture medium was refreshed every 3 days. At the third passage, the cells were ready for EVs collection.

### Isolation of Extracellular Vesicles

Cells were grown until they reached 90–95% confluency. The growth medium was then aspirated, and the cells were rinsed with DPBS before being detached using a 0.25% trypsin solution. To neutralize the trypsin, DMEM containing 10% FBS was added. Following this, the cells were centrifuged at 1400 rpm for 5 minutes. To remove any residual serum, the cells were washed with 0.9% NaCl. Next, the cells were incubated for 30 minutes in serum-free DMEM supplemented with 10 μg/mL cytochalasin B (Sigma-Aldrich, Burlington, MA, USA) at 37°C and 5% CO2. After incubation, the cell suspension was vigorously mixed on a vortex for 60 seconds and then subjected to centrifugation at 500 rpm for 10 minutes. The supernatant was collected and further centrifuged at 700 rpm for 10 minutes, followed by a final centrifugation at 12,000 rpm for 15 minutes. The resulting pellet, containing EVs, was resuspended in 0.9% NaCl.

### Spinal Cord Injury and MSC-EVs Therapy

The study was carried out on adult female Wistar rats (n=23) weighing 250–300 g (Pushchino Laboratory, Russia). The animals were housed in a 12-hour light/dark cycle conditions with food and water available *ad libitum*. Anesthesia was administered using isoflurane (1.3%) and Zoletil (20 mg/kg) for all surgical procedures. Following a laminectomy, a moderate spinal cord contusion injury (SCI) was induced at the Th8 level with an impact speed of 2.5 m/s using the Impact One Stereotaxic Impactor (Leica). The animals were divided into four experimental groups. In the first two experimental groups, 5 and 10 µg of MSC-EVs encapsulated in 18 µl of fibrin matrix (FM, Baxter, USA) were applied immediately after injury (SCI FM+EVs5, n=6 and SCI FM+EVs10, n=5, groups, respectively). In the first control group, animals did not receive MSC-EVs therapy (SCI group, n=6), while in the second control group, FM without MSC-EVs was applied immediately after injury (SCI FM group, n=6). After surgery, all experimental rats received daily intramuscular doses of gentamicin (25 mg/kg, Microgen, Russia) for 7 days. Bladders of the injured rats were manually emptied twice daily until spontaneous urination was restored.

### Multiplex Assay

To assess the cytokine profile of the injured spinal cord at 60 days post-injury (dpi) a section of the spinal cord, including epicenter of injury, was dissected at the Th8 level. The tissue was homogenized using an electric homogenizer with 300 µl of complete extraction buffer. Following centrifugation at 13,000 rpm for 20 minutes at 4°C, the soluble protein extract was collected and stored at −80°C until cytokine analysis. The cytokine profile was determined using multiplex analysis with xMAP Luminex technology, specifically the Bio-Plex Pro Rat Cytokine 23-Plex Immunoassay (12005641, Bio-Rad, USA). This assay allows for the quantification of 23 cytokines and chemokines, including G-CSF, GM-CSF, GRO/KC, IFN-γ, IL-1α, IL-1β, IL-2, IL-4, IL-5, IL-6, IL-7, IL-10, IL-12 (p70), IL-13, IL-17A, IL-18, M-CSF, MCP-1, MIP-1α, MIP-3α, RANTES, TNF-α, and VEGF.

### Statistical analysis

The obtained results were processed using the Origin Pro software. Data are presented as mean values with standard deviation (SD) or standard error (SE). A normality test was conducted for all study groups. One-way analysis of variance (ANOVA) followed by Tukey’s test was performed for multiple group comparisons. All analyses were presented in a blinded manner relative to the study groups. Significant differences below the value of 0.05 (P<0.05) were applied for all statistics.

## Results

### Assessment of cytokine profile changes

The analysis of cytokine levels in spinal cord homogenates from the experimental groups (SCI, SCI FM, SCI FM+EVs5, and SCI FM+EVs10) 60 dpi revealed notable differences in the expression of the cytokines studied. The data obtained through multiplex analysis (Supplementary Table 1) were visualized as a heat map (Figure 2).

**Figure 2.**
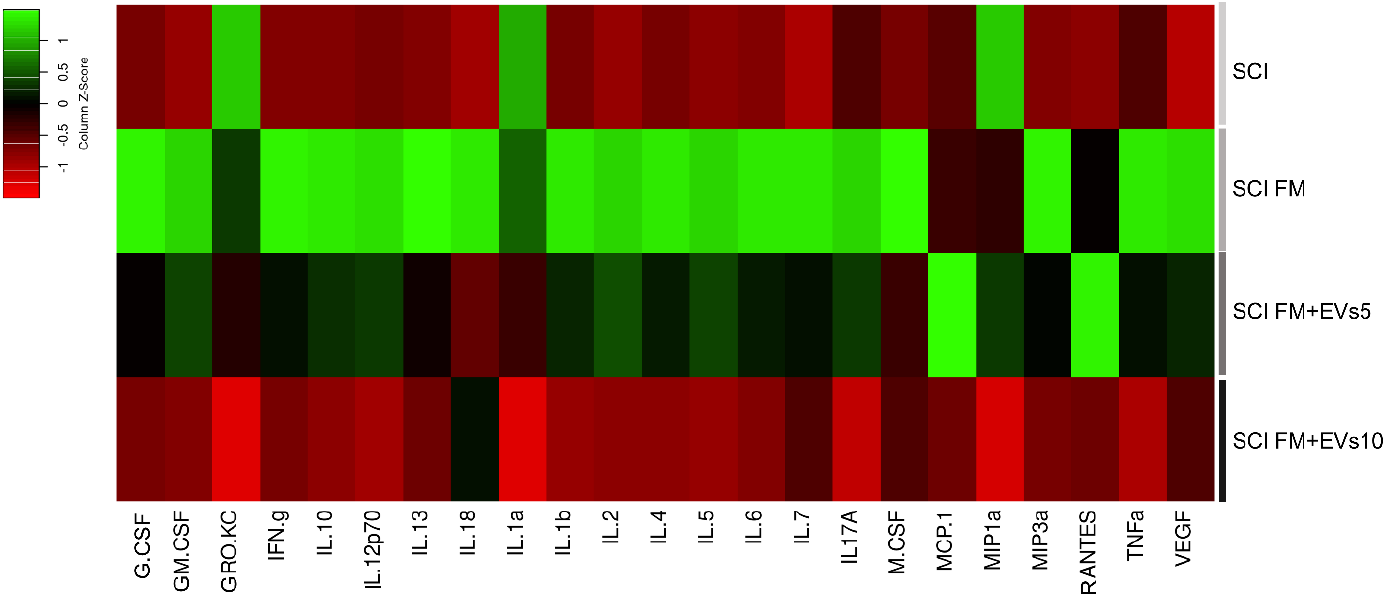
Multiplex analysis of spinal cord homogenates (Th8) from control (SCI, SCI FM) and experimental (SCI FM+EVs5, SCI FM+EVs10) groups. A heat map shows cytokines upregulated (green) or downregulated (red).

The expression levels of several cytokines were significantly elevated in the second control group (SCI FM) compared to the first control group (SCI). Notably, G-CSF showed the highest increase, with a 2.8-fold elevation (P <0.05) (Figure 3A). Other cytokines also exhibited significant increases, including GM-CSF (1.6-fold), IFN-γ (2.3-fold), IL-18 (1.2-fold), IL-2 (2-fold), IL-6 and IL-7 (both 2.4-fold), M-CSF (1.7-fold), and MIP3a (2.3-fold).

**Figure 3.**
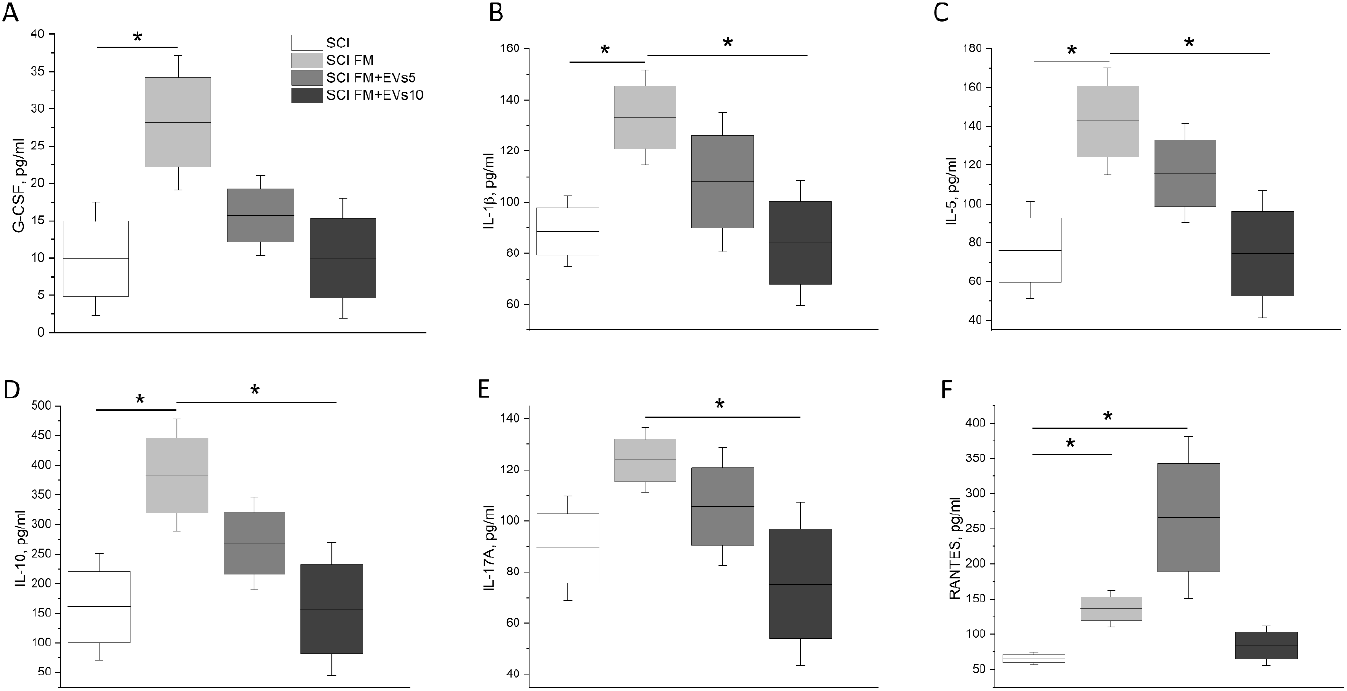
Multiplex analysis. Boxplots display the concentrations (Y-axis) of six cytokines: (A) G-CSF, (B) IL-10, (C) IL-1β, (D) IL-5, (E) IL-17A, and (F) RANTES across four experimental groups: SCI (spinal cord injury without treatment), SCI FM (fibrin matrix only), SCI FM+EVs5 (5 µg EVs in fibrin matrix), and SCI FM+EVs10 (10 µg EVs in fibrin matrix). *P<0.05, ANOVA with Tukey’s post hoc test.

For cytokines IL-1β, IL-5, and IL-10, the mean concentrations in spinal cord homogenates of the second control group (SCI FM) showed a significant increase compared not only to the SCI group but also to the SCI FM+EVs10 group (P<0.05) (Figure 3B-D). The application of a 10 µg dose of EVs (SCI FM+EVs10) to the lesion area significantly reduced the concentrations of both pro-inflammatory cytokines, IL-1β (1.5-fold) and IL-5 (1.9-fold), as well as the anti-inflammatory cytokine IL-10 (2.4-fold).

A similar trend was observed for IL-17A expression, where a 1.6-fold decrease (P < 0.05) was found in the SCI FM+EVs10 group compared to the SCI FM control group (Figure 3E). Notably, the results for RANTES expression (Figure 3F) should be highlighted, as the level in the SCI FM+EVs5 group was significantly higher than in the SCI group (a 4-fold increase, P < 0.05). Additionally, there were significant differences in RANTES levels between the two control groups (SCI FM vs. SCI), with a 2-fold higher expression in the SCI FM group (P < 0.05).

## Discussion

In this study, we conducted a multiplex analysis of 23 cytokines in rat spinal cord homogenates during the chronic phase of SCI (60 dpi) following the application of FM combined with MSC-EVs therapy at two dosages (5 µg and 10 µg). The results were compared to SCI without treatment and SCI treated with FM alone. Our analysis revealed an upregulation of several cytokines following MSC-EVs therapy.

In the case of FM application alone, we observed a tendency toward a sustained increase in chemokines (G-CSF, M-CSF, GM-CSF) and most of the cytokines studied within the SCI region, particularly those with predominantly pro-inflammatory effects (IL-1β, IL-2, IL-5, IL-6, IL-7, IL-17A, IL-18, IFN-γ, and MIP-3α), alongside a more moderate rise in anti-inflammatory cytokines (IL-10). Previous studies indicate that fibrin glues, such as Tisseel® (also referred to as Tissucol® in our study), despite their hemostatic properties, can elicit local inflammatory responses [7,8]. In one study, no attenuation of inflammation was noted when fibrin glue was used in a rat SCI model, which may be linked to its effects on coagulation processes [7]. These properties of fibrin glue might activate inflammatory cascades, which could account for the elevated levels of both pro-inflammatory and anti-inflammatory cytokines in our experiment. Additionally, a study comparing Tisseel® with other matrixes, such as BioGlue® and Adherus®, demonstrated that Tisseel® caused relatively less pronounced inflammatory and degenerative responses in a rat SCI model [8]. However, even in these cases, localized inflammatory reactions were still observed, aligning with our findings. It is noteworthy that these studies examined the effects of adhesives on inflammation up to 28 days post-application, whereas our study extended to 60 dpi, suggesting that the inflammatory responses may be prolonged.

The study of FM biodegradation in the body is also noteworthy. As a biocompatible material, Tisseel® is slowly resorbed in tissues, which may lead to prolonged interactions with surrounding cells, potentially sustaining an inflammatory response. Research on various commercial fibrin matrices, including Tisseel®, has shown that the presence of protease inhibitors like aprotinin slows its biodegradation, suggesting that its extended presence in tissues may have prolonged effects on cellular responses [9]. In an *in vitro* study, Tisseel® was also found to increase levels of metalloproteinases (MMP-1, MMP-2) in mesothelial cells and fibroblasts, which may result in altered cytokine profiles and the persistence of inflammation during matrix degradation [10]. Furthermore, studies have identified key cellular targets and molecular mechanisms by which fibrinogen and thrombin, essential components of Tisseel®, can hinder neurotrauma recovery. For instance, fibrinogen binding to αVβ3 inhibits neurite outgrowth in the CNS, as demonstrated in a mouse model of encephalomyelitis [11], while its interaction with CD11b/CD18 activates microglia in a mouse model of traumatic brain injury [12]. These mechanisms could potentially interfere with the natural tissue repair processes following traumatic injuries. It is also important to note that differences in viral inactivation and processing methods between Tisseel® and Vistaseal® during production have been suggested to theoretically influence the inflammatory potential of Tisseel®. However, the clinical significance of these differences has not yet been fully established [13].

Our study is among the first to explore the effects of FM combined with MSC-EVs on the cytokine profile during the chronic phase of SCI (60 dpi). While previous research has examined the role of MSC-EVs encapsulated in FM in promoting oligodendrogenesis and functional recovery [14], as well as neurogenesis [15] in rodent models of chronic phase of SCI, the impact of this combination on neuroinflammatory processes within the injured spinal cord remains unstudied. We found that MSC-EVs can mitigate cytokine shifts at the injury site induced by local FM exposure, exhibiting a clear dose-dependent effect. Specifically, increasing the MSC-EVs dosage significantly reduced the expression levels of cytokines IL-1β, IL-5, IL-17A, and IL-10 compared to the FM-only group. However, the notably elevated levels of the chemokine RANTES (CCL5) observed with the 5 µg EV dose, as opposed to 10 µg, warrants further investigation to clarify its role in anti-inflammatory or regenerative processes.

MSC-EVs have been shown to significantly inhibit the activation of the NLRP3 inflammasome and p38/MAPK signaling pathways, both of which are critical in triggering pro-inflammatory responses in traumatic brain injury models in mice [16]. This inhibition led to reduced production of pro-inflammatory cytokines, including IL-1β and IL-6, mediating an overall immunomodulatory effect [17]. Furthermore, animals treated with MSC-EVs exhibited improved cognitive and motor functions, reduced long-term inflammation, and decreased brain damage [16,18]. In a separate study involving a rodent model of SCI, administration of MSC-EVs was associated with a decrease in pro-inflammatory cytokines (IL-6, IL-1β), suppression of microglial reactivity [19,20], and anti-apoptotic activity, leading to reduced neuroinflammation and enhanced recovery [21]. Additionally, MSC-EVs administration resulted in a significant reduction in TNF-α and IL-1β levels, activating autophagy, which further contributed to inflammation reduction and tissue regeneration [22]. Thus, our findings align with previous studies demonstrating that MSC-EVs modulate inflammatory processes by lowering pro-inflammatory cytokine levels and suppressing microglial activity in traumatic CNS injury models.

## Conclusion

Our study demonstrated that the fibrin matrix itself plays a crucial role in influencing inflammatory processes, as it promotes both pro-inflammatory and anti-inflammatory responses. This likely occurs through the activation of cells involved in tissue repair. Additionally, the application of MSC-EVs, encapsulated in the fibrin matrix, significantly modulates the levels of both pro-inflammatory and anti-inflammatory cytokines in the chronic phase of SCI. The dose-dependent effects observed, particularly with the 10 µg MSC-EVs dose, suggest that this combination therapy could offer a novel approach for modulating inflammatory responses and promoting tissue regeneration post-injury.

## Authors’ contributions

Conceptualization, Y.M.; methodology, Y.M., I.K.; software, A.R.; validation, E.G., Y.M., I.K.; formal analysis, E.G., A.K., I.K.; investigation, A.K., I.K., E.G.; resources, Y.M., A.R.; data curation, Y.M.; writing—original draft preparation, Y.M., A.K., I.K.; writing—review and editing, Y.M., I.K.; visualization, I.K.; supervision, Y.M.; project administration, Y.M., A.R.; funding acquisition, Y.M., A.R. All authors have read and agreed to the published version of the manuscript.

## Availability of data and materials

The data presented in this study are available on request from the corresponding author. The data are not publicly available due to the evolving nature of the project.

## Ethical approval and consent to participate

The methods described herein were performed in accordance with the Declaration of Helsinki and approved by the local ethical committee of Kazan (Volga region) Federal University (No. 2, May 5, 2015).

## Consent for publication

Not applicable.

