## Supplementary Table 1 for "Local cytokine changes in response to mesenchymal stem cell-derived extracellular vesicles-based therapy in rat spinal cord injury"

**Supplementary Table 1**. Multiplex analysis of spinal cord homogenates (Th8) from control (SCI, SCI FM) and experimental (SCI FM+EVs5, SCI FM+EVs10) groups (pg/ml).

| **SCI** | **G-SCF** | | **GM-CSF** | **GRO/K** | **IFN-g** | **IL-10** | **IL-12p70** | **IL-13** | **IL-18** | **IL-1a** | **IL-1b** | **IL-2** |
| --- | --- | --- | --- | --- | --- | --- | --- | --- | --- | --- | --- | --- |
| Mean value | 9,90 | | 36,65 | 46,15 | 198,42 | 160,71 | 67,52 | 83,6 | 968,96 | 127,2 | 88,45 | 491,42 |
| Standard deviation | 11,37 | | 18,20 | 17,68 | 167,21 | 147,75 | 45,12 | 67,25 | 154,43 | 45,73 | 22,71 | 236,89 |
| **SCI** | **IL-4** | **IL-5** | **IL-6** | **IL-7** | **IL17A** | **M-CSF** | **MCP-1** | **MIP1a** | **MIP3a** | **RANTES**^#^ | **TNFa** | **VEGF** |
| Mean value | 29,54 | 76,15 | 209,64 | 20,02 | 89,39 | 40,22 | 417,28 | 76,32 | 10,02 | 65,4 | 382,69 | 58,39 |
| Standard deviation | 26,89 | 41,12 | 207,07 | 16,00 | 33,37 | 21,02 | 145,71 | 28,41 | 7,21 | 14,97 | 72,71 | 37,16 |

| **SCI FM** | **G-SCF*** | | **GM-CSF*** | **GRO/KC** | **IFN-g*** | **IL-10**** | **IL-12p70** | **IL-13** | **IL-18*** | **IL-1a** | **IL-1b**** | **IL-2*** |
| --- | --- | --- | --- | --- | --- | --- | --- | --- | --- | --- | --- | --- |
| Mean value | 28,17 | | 60,38 | 41,97 | 454,60 | 383,05 | 122,72 | 173,94 | 1229,89 | 115,34 | 133,09 | 955,11 |
| Standard deviation | 14,70 | | 15,97 | 11,67 | 182,50 | 155,00 | 43,9 | 67,58 | 214,61 | 18,01 | 30,48 | 368,14 |
| **SCI FM** | **IL-4** | **IL-5**** | **IL-6*** | **IL-7*** | **IL17A**** | **M-CSF*** | **MCP-1** | **MIP1a** | **MIP3a*** | **RANTES** | **TNFa** | **VEGF** |
| Mean value | 68,09 | 142,57 | 504,43 | 48,88 | 123,77 | 68,17 | 425,51 | 65,68 | 23,32 | 136,11 | 443,78 | 91,38 |
| Standard deviation | 31,18 | 44,97 | 214,77 | 18,91 | 20,43 | 19,09 | 160,03 | 31,63 | 9,43 | 41,6 | 66,61 | 41,92 |

| **SCI FM+EVs5** | **G-SCF** | | **GM-CSF** | **GRO/KC** | **IFN-g** | **IL-10** | **IL-12p70** | **IL-13** | **IL-18** | **IL-1a** | **IL-1b** | **IL-2** |
| --- | --- | --- | --- | --- | --- | --- | --- | --- | --- | --- | --- | --- |
| Mean value | 15,73 | | 51,06 | 39,15 | 299,97 | 267,65 | 96,98 | 112,00 | 1012,03 | 92,7 | 107,86 | 783,93 |
| Standard deviation | 8,8 | | 16,61 | 14,59 | 161,49 | 129,06 | 56,27 | 59,22 | 387,98 | 44,66 | 44,21 | 368,62 |
| **SCI FM+EVs5** | **IL-4** | **IL-5** | **IL-6** | **IL-7** | **IL17A** | **M-CSF** | **MCP-1** | **MIP1a** | **MIP3a** | **RANTES** | **TNFa** | **VEGF** |
| Mean value | 44,95 | 115,92 | 337,51 | 33,44 | 105,64 | 44,9 | 497,4 | 70,47 | 14,91 | 266,24 | 401,07 | 76,27 |
| Standard deviation | 20,64 | 41,8 | 194,33 | 14,30 | 37,57 | 21,6 | 51,2 | 33,13 | 6,51 | 187,73 | 155,25 | 29,55 |

| **SCI FM+EVs10** | **G-SCF** | | **GM-CSF** | **GRO/K** | **IFN-g** | **IL-10** | **IL-12p70** | **IL-13** | **IL-18** | **IL-1a** | **IL-1b** | **IL-2** |
| --- | --- | --- | --- | --- | --- | --- | --- | --- | --- | --- | --- | --- |
| Mean value | 9,99 | | 38,09 | 33,75 | 204,49 | 156,77 | 61,94 | 89,95 | 1085,37 | 67,43 | 84,05 | 510,74 |
| Standard deviation | 11,97 | | 26,77 | 11,93 | 200,25 | 168,55 | 55,11 | 80,9 | 193,01 | 49,62 | 32,49 | 322,05 |
| **SCI FM+EVs10** | **IL-4** | **IL-5** | **IL-6** | **IL-7** | **IL17A** | **M-CSF** | **MCP-1** | **MIP1a** | **MIP3a** | **RANTES** | **TNFa** | **VEGF** |
| Mean value | 27,98 | 74,14 | 214,9 | 26,89 | 75,34 | 43,21 | 413,48 | 58,95 | 10,27 | 83,79 | 363,19 | 67,54 |
| Standard deviation | 28,69 | 48,99 | 215,75 | 14,42 | 47,59 | 33,27 | 73,81 | 34,64 | 8,65 | 42,15 | 136,28 | 57,41 |

* - P<0.05 compared to SCI group; ** - P<0.05 compared to SCI and SCI FM+EVs10 groups; ^#^ - P<0.05 compared to SCI FM and SCI FM+EVs5 groups
